# Immediate early gene activation throughout the brain is associated with dynamic changes in social context

**DOI:** 10.1101/275495

**Authors:** Cait M. Williamson, Inbal S. Klein, Won Lee, James P. Curley

## Abstract

Social competence is dependent on successful processing of social context information. The social opportunity paradigm is a methodology in which dynamic shifts in social context are induced through removal of the alpha male in a dominance hierarchy, leading to rapid ascent in the hierarchy of the beta male and of other subordinate males in the social group. In the current study, we use the social opportunity paradigm to determine what brain regions respond to this dynamic change in social context, allowing an individual to recognize the absence of the alpha male and subsequently perform status-appropriate social behaviors. Replicating our previous work, we show that following removal of the alpha male, beta males rapidly ascend the social hierarchy and attain dominant status by increasing aggression towards more subordinate individuals. Analysis of patterns of Fos immunoreactivity throughout the brain indicates that in individuals undergoing social ascent, there is increased activity in regions of the social behavior network, as well as the infralimbic and prelimbic regions of the prefrontal cortex and areas of the hippocampus. Our findings demonstrate that male mice are able to respond to changes in social context and provide insight into the how the brain processes these complex behavioral changes.

## INTRODUCTION

Organization into dominance hierarchies is a fundamental feature of group social behavior across species, including non-human primates (Muller & Wrangham, 2004; Sapolsky, 1993), cichlid fish (Grosenick, Clement, & Fernald, 2007; Huffman, Hinz, Wojcik, Aubin-Horth, & Hofmann, 2015; Oliveira & Almada, 1996), naked mole rats (Holmes, Goldman, & Forger, 2008), honey bees (Kucharski, Maleszka, Foret, & Maleszka, 2008), mice (Wang et al., 2011; Williamson, Lee, & Curley, 2016a), and humans (Zink et al., 2008). Individuals form these dominance structures through a complicated appraisal of their social context in order to ascertain their position relative to that of the other individuals within their social network (Curley, 2016b; Fernald, 2014; Grosenick et al., 2007; Oliveira, 2009). There has been characterization of the complex behavioral features of the formation and maintenance of dominance hierarchies (Chase & Seitz, 2011; Chase, Tovey, Spangler-Martin, & Manfredonia, 2002; Curley, 2016b; Williamson, Lee, et al., 2016a; Williamson, Lee, Romeo, & Curley, 2017), as well as identification of the neural correlates associated with social status in stable social hierarchies (So, Franks, Lim, & Curley, 2015; Wang et al., 2011; Williamson, Franks, & Curley, 2016; Zerubavel, Bearman, Weber, & Ochsner, 2015; Zink et al., 2008).

Although hierarchies are commonly stable, there often occurs times when individuals change in social rank. One particularly salient example of this is when a power vacuum emerges at the top of a hierarchy following the removal or deposition of the alpha individual. When such social opportunities occur, subdominant animals typically rapidly ascend to the alpha position. Such behavior has been observed experimentally in hierarchies of both cichlid fish (Maruska & Fernald, 2010) and CD1 outbred mice (Williamson, Romeo, & Curley, 2017) associated with changes along the hypothalamic-pituitary-gonadal (HPG) axis. Ascent to dominant status in cichlid fish is also associated with increased immediate early gene expression in several regions specific to fish social behavior (Burmeister, Jarvis, & Fernald, 2005; Maruska, Zhang, Neboori, & Fernald, 2013). However, there has been no comprehensive, whole brain analysis of the neural response to changes in social context in mammals. This ability to process this dynamic social context information and behave in a socially competent manner when the structure of a social hierarchy shifts is critical for successful social living.

The “Social Behavior Network” (SBN) is a bidirectional circuit of brain regions associated with multiple forms of social behavior (i.e. aggression, sexual behavior, communication, social recognition, affiliation and bonding, parental behavior, and social stress responses) across species (Goodson, 2005; Newman, 1999). This network was first described to include the medial amygdala (meA), the bed nucleus of the stria terminalis (BNST), the lateral septum (LS), the medial preoptic area (mPOA), the anterior hypothalamus (AH), the ventromedial hypothalamus (VMH), and the periaqueductal grey (PAG) (Newman, 1999). These brain regions are thought to be the core of the social brain, with much supporting evidence for their role in regulating relatively simple social behavior (see Goodson, 2005 for a comprehensive review). However, for complex social behaviors, such as the formation, maintenance, and dynamic adjustment of social hierarchies, which are reliant on an individual’s ability to perceive changes in their social environment, it is important to understand how activity within the SBN is modulated and complemented by brain regions associated with executive functioning (i.e. prefrontal cortex (Wang et al., 2011; Zink et al., 2008)) and memory (i.e. hippocampus (Noonan et al., 2014; Williamson, Franks, et al., 2016)).

In previous studies, we have demonstrated differential gene expression throughout the brains of outbred CD1 mice of different social rank living in linear hierarchies, specifically in the medial amygdala, central amygdala, medial preoptic area (So et al., 2015) and in the whole hippocampus (Williamson, Franks, et al., 2016). We have shown that within minutes of the removal of the dominant male from a social group, the subdominant male exhibits increased aggression as well as rapid changes in GnRH gene expression in the medial preoptic region of the hypothalamus (Williamson, Romeo, et al., 2017). In the current study, we aimed to generate a map of immediate early gene activity throughout the SBN and areas related to the monitoring of social context and social memory to assess how the brain of subdominant animals responds to a changing social context when a social opportunity to ascertain alpha status arises. Specifically, we assessed the pattern of Fos immunoreactivity in subdominant mice in response to the removal of the alpha male (a dynamic social change) and compared this neural response to that of subdominant mice living in a stable social system.

## METHODS

### Subjects and Housing

Throughout the study, mice were housed in the animal facility in the Department of Psychology at Columbia University, with constant temperature (21–24°C) and humidity (30-50%) and a 12/12 light/dark cycle with white light (light cycle) on at 2400 hours and red light (dark cycle) on at 1200 hours. All mice were individually and uniquely marked by dying their fur with a blue, nontoxic, non-hazardous animal marker (Stoelting Co.). These marks remain for up to 12 weeks and only require one application, enabling each animal to be visually identified throughout the study. All procedures were conducted with approval from the Columbia University Institutional Animal Care and Use Committee (IACUC – Protocol No: AC-AAAP5405).

### Behavioral Manipulation

To determine Fos activation associated with social ascent, we performed a social opportunity manipulation comparing subdominant individuals in socially stable groups to those ascending in a hierarchy from subdominant to dominant status. This procedure is similar to that previously described (Williamson, Romeo, & Curley, 2017). A total of 48 male outbred CD1 mice aged 7 weeks were obtained from Charles River Laboratories and housed in groups of 3 for 2 weeks in standard sized cages. At 9 weeks of age, groups of 12 mice were placed into custom built vivaria (length 150cm, height 80cm, width 80cm; Mid-Atlantic; **Supplemental Figure 1**). The vivarium was constructed as previously described (Williamson, Lee, & Curley, 2016). Each vivarium consists of an upper level consisting of multiple shelves covered in pine bedding and a lower level consisting of a series of nest-boxes filled with pine bedding connected by tubes. Mice can access all levels of the vivarium via a system of ramps and tunnels. Standard chow and water were provided ad libitum at the top of the vivarium, encouraging movement and exploration of all the shelves. Social groups (N = 12 per group) were paired such that each group was introduced into the vivarium directly before onset of the dark cycle on the same day as one other group, creating 2 sets of paired cohorts (4 cohorts total). Live behavioral observations were conducted each day during the dark cycle. These observations consisted of trained observers recording all instances of fighting, chasing, mounting, subordinate posture, and induced-flee behaviors. The identity of the dominant and subordinate individuals in each interaction were recorded using all occurrence sampling. Data was collected directly into electronic tablets and uploaded live to a google spreadsheet. **Supplemental Table 1** contains an ethogram of the behaviors recorded. Observations were conducted for 2 hours each day on Days 1-4 of group housing during the dark (red) light cycle. At the end of Day 4, it was verified that a dominance hierarchy had emerged in each group, and the identity of the alpha and beta male in each group was determined. On Day 5, at the onset of the dark cycle, the alpha male from one of the paired cohorts was removed from the vivarium and placed in a standard cage with food and water. In the other paired cohort, the alpha male was sham-removed, which entailed an experimenter opening the Perspex windows to the vivarium, placing their hand in the vivarium, and reaching towards the alpha mouse but not removing it from the vivarium. This condition, which does not involve removing any mouse from the social group, controls for behavioral changes that may be occurring in response to a non-social disturbance to the environment. Live behavioral observations occurred for the period directly following the removal or sham-removal. Ascending males were confirmed as the individual who won most aggressive contests post-alpha removal without consistently losing to other individuals. Ninety minutes after this ascending individual had won three fights, the ascending male was removed from the alpha removal group and the non-ascending subdominant male was removed from the sham-removal group.

Following removal from the social group, mice were anesthetized with ketamine/xylazine and perfused intracardially with phosphate-buffered saline (PBS) followed by 4% paraformaldehyde. Brains were stored at 4°C in 4% paraformaldehyde for the first six hours following perfusion and then switched to a 30% sucrose solution. Following the perfusions, the alpha male who had been removed in the social ascent condition was returned to his social group. Alpha males always retained their alpha status on return to the social group.

This procedure was repeated at four day intervals for a total of six “removals”. Manipulations were counter-balanced between paired cohorts (i.e. one vivarium had alpha removal for removals 1, 3, and 5, and sham removals for removals 2, 4, and 6, and the opposite was true for the paired vivarium). Each removal/sham-removal decreased the size of the social group by 1, resulting in N = 12 (removal one), N = 11 (removal two), N = 10 (removal three), N = 9 (removal 4), N = 8 (removal 5), N = 7 (removal 6). This design yielded 12 mice per group from two groups: beta male/alpha removed (ascending males) and beta male/alpha remained (sham-removal subdominants). See **Figure 1** for a schematic of the behavioral manipulation.

**Figure 1.**
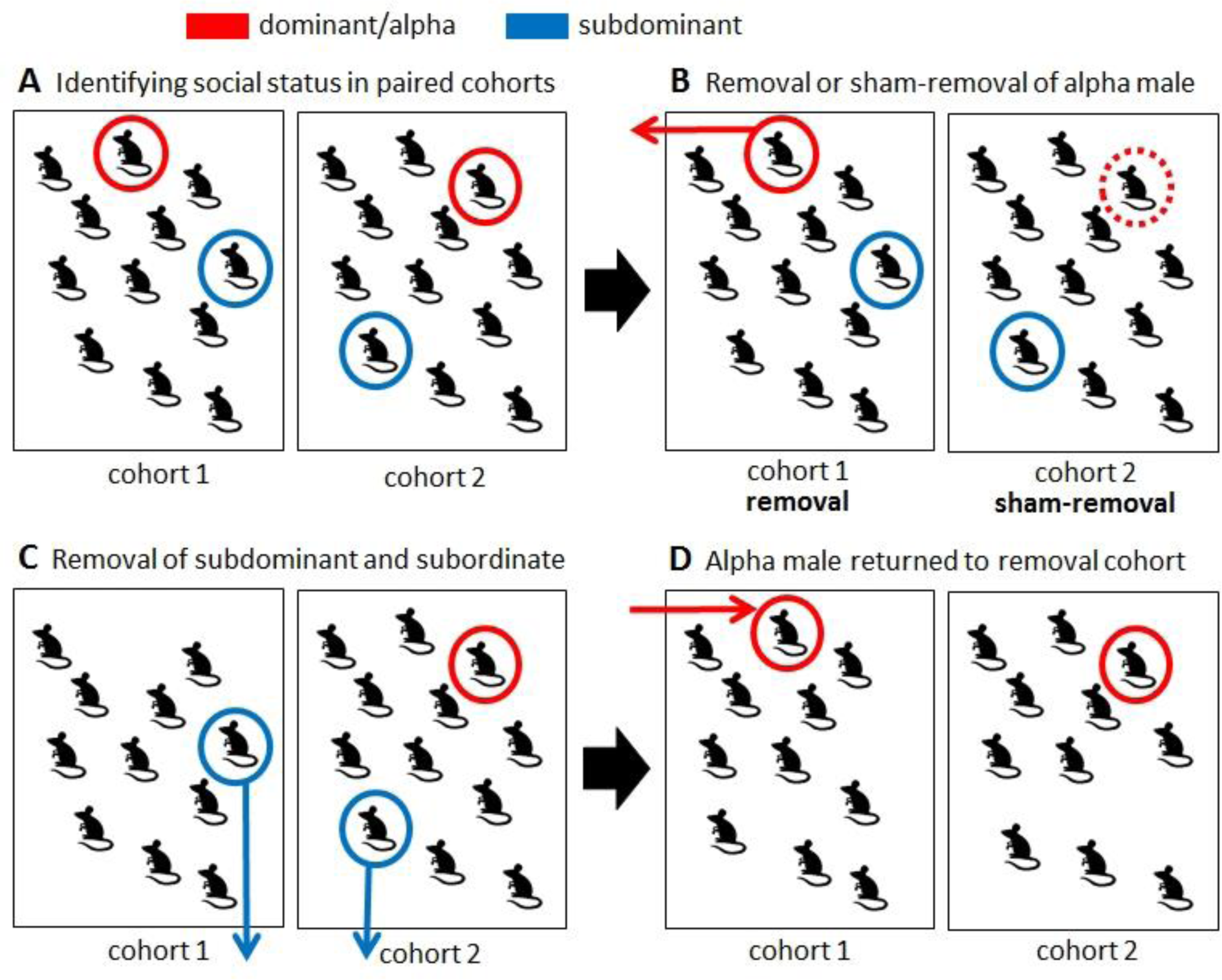
Schematic of the social opportunity experimental design. (A) Two cohorts of twelve mice are put into separate vivaria and a stable social hierarchy emerges, with clearly defined dominant, and subdominant individuals. (B) The alpha male is removed from one stable hierarchy and sham-removed from the paired hierarchy. (C) Following removal/sham-removal, behavioral observations are conducted on both cohorts until ninety minutes after a subdominant rises in the alpha-removed group. At this ninety-minute time point, the sub-dominant (rising to alpha) animal in each hierarchy is removed and brain extracted for analyses. (D) One-hour following this removal of the subdominant, the alpha male is returned to its social group. This procedure is repeated five more times four days apart for each pair of cohorts.

### Immunohistochemistry

Brains were stored in 30% sucrose in 0.1M PB at 4°C until slicing. Perfused whole brains were sliced coronally into 40 µm sections and stored in 0.1 M PB azide until processing according to the avidin– biotin procedure, using the Vectastain ABC Elite peroxidase rabbit IgG kit (Vector Laboratories, Burlingame, CA). Free-floating sections were transferred into wells and washed three times in 0.1 M PB for five minutes each rinse. The sections were then washed once in hydrogen peroxide for five minutes and then washed three times in PBT for five minutes each rinse. The sections were then placed in a solution of 2% Normal Goat Serum (NGS, Vector Laboratories) in 0.1% Triton-X in 0.1M PB (PBT) for an hour, and then incubated in primary Fos rabbit polyclonal IgG (Santa Cruz, USA, SC-52) at a concentration of 1:5000 overnight at 4˚C with 2% NGS block. The next day, the sections were washed 3 times in PBT for 5 minutes each rinse and then incubated in biotinylated anti-rabbit IgG (Vecstastain ABC Kit, Vector Laboratories) at a concentration of 1:200 in PBS for 1 hour. Once the hour was complete, sections were once again washed 3 times in PBT for 5 minutes each rinse. Sections were then incubated for 1 hour in an avidin– biotin–peroxidase complex in 0.1 M PBT (A and B solutions of the Vectastain ABC Kit, Vector Laboratories) at a concentration of 40ul A: 40ul B: 10ml PBT and then washed 3 times in 0.1M PBS for 5 minutes each rinse. Fos immunoreactivity was visualized by incubating the sections in 0.02% 3,3’-diaminobenzidine (DAB) solution for 2-4 minutes. Fos immunoreactivity was visualized through incubation in a DAB stain for 2-4 minutes. Sections were then washed once for 1 minute in 0.1M PBS and then washed 3 times in 0.1M PBS for 5 minutes each rinse. All sections were then stored in 0.1M PB at 4˚C for up to 24 hours until mounting. Sections were mounted onto FisherBrand Plus slides and then coverslipped with DePeX mounting medium (Sigma-Aldrich, St. Louis, MO).

### Photos and Image Analysis

Images were taken of brain sections under a 10x objective microscope at a magnification x100 and a digital camera. Localization of specific brain regions was determined using the Allen Mouse Brain Atlas (Lein et al., 2007). Images were then cropped to include only the exact portion of each brain region by overlaying images from *The Mouse Brain in Stereotaxic Coordinates* (Paxinos & Franklin, 2004) over the photos in an image editing program. Particles were then analyzed with the batch function using a macro in ImageJ (Schneider, Rasband, & Eliceiri, 2012). Twenty-five separate brain regions were processed: bed nucleus of the stria terminalis (BNST), lateral septum (LS), anterior hypothalamus (AH), medial preoptic area (mPOA), ventromedial hypothalamus (VMH), medial amygdala (meA), dorsolateral periaqueductal grey (dlPAG), ventrolateral periaqueductal grey (vlPAG), dorsal and ventral premammillary nuclei (PMd and PMv), cingulate cortex, infralimbic and prelimbic regions of the prefrontal cortex (IL, PrL), piriform cortex, retrosplenial cortex (RC), area CA1 of the hippocampus (CA1), area CA3 of the hippocampus (CA3), the dentate gyrus (DG), anterior cortical amygdala (ACA), central amygdala (CeA), basolateral amygdala (BLA), arcuate nucleus (Arc), lateral hypothalamus (LH), primary auditory cortex, primary visual cortex. All subjects for each brain region were analyzed concurrently, with each brain region being analyzed separately.

### Statistical Analysis

All statistical analyses were undertaken in R version 3.4.0 (R Core Team, 2016) in RStudio version 1.0.143 (RStudio Team, 2015).

#### Behavioral analysis

For each cohort, the linearity of the social hierarchy was calculated using Landau’s Modified h’. Briefly, the total number of wins by each individual against all other individuals are entered into a sociomatrix. Landau’s method then assesses the degree to which each individual consistently dominates others in contests and whether individuals can be linearly ordered based upon their wins and losses. The h’ value ranges from 0 (no linearity) to 1 (completely linear). The significance of h’ is determined by performing 10,000 two-step Monte Carlo randomizations of the sociomatrix and comparing the observed h’ against a simulated distribution of h’ (De Vries, 1995; Williamson, Lee, & Curley, 2016b). Temporal changes in individual dominance ratings were calculated using Glicko Ratings (Glickman, 1999; So et al., 2015). Glicko ratings are a pairwise-contest model ratings system where ratings points are recalculated following each successive win or loss. All individuals start with a rating of 2200. Ratings are gained after wins and lost after losses with the magnitude of points gained or lost dependent upon the difference in ratings scores between the two individuals in each contest (Glickman, 1999; Williamson, Lee, et al., 2016b). Landau’s modified h’ was calculated using the R package compete v0.1 (Curley, 2016a). Glicko ratings were calculated using the PlayerRatings package v1.0 in R (Stephenson & Sonas, 2012).

#### Social ascent analysis

To compare wins and losses between betas in the alpha removed group to those in the alpha remained group, we used Wilcoxon rank sum tests. To compare wins and losses between betas in the alpha removed and sham-removed group on the day of removal or sham-removal to their behavior the day before, we used Wilcoxon signed rank tests.

#### Fos Analysis

To determine the effect of alpha removal on the number of immunoreactive cells in each brain region, we used the R package lme4 (Bates et al., 2015) to run negative binomial mixed models, with social status (alpha removed or alpha remained) as a fixed effect and cohort, removal number, side of the brain, number of wins, and number of losses as random effects.

#### Hierarchical clustering analysis

To determine brain region activation patterns in both the alpha removed and alpha remained groups, we created correlation matrices for each group and visualized them using the R package lattice (Sarkar, D, 2017). We then used the package pvclust (Suzuki & Simodaira, 2015) to determine hierarchical clusters and generate p-values for each cluster using multiscale bootstrap resampling.

## RESULTS

### All cohorts form significantly linear hierarchies

All social groups formed significantly linear dominance hierarchies with a clear alpha and beta male after the first four days of group housing prior to the first alpha or sham-removal (all h’ values > 0.45, mean h’ = 0.59, all p < 0.05, mean p = 0.016). All alpha males maintained their alpha status for the duration of their presence in the established social hierarchy.

### Subdominant males socially ascend following removal of the dominant male

After each of the 12 alpha male removals, a subdominant male ascended within 1 hour. Rising subdominants had significantly more wins than the subdominant males in the sham-removal group (Wilcoxon rank sum test W = 138, p = 0.00018 – **Figure 2A**). Further, the majority (9/12) of rising subdominant individuals never lost a fight during this period, however there was no significant difference in number of losses between rising subdominants in the alpha removed group and those in the sham removal group (Wilcoxon rank sum test W = 56.5, p = 0.3006 – **Figure 2B**). When compared to their behavior the day before alpha removal, rising subdominants had significantly more wins (Wilcox signed rank test V = 1, p 0.003) and significantly fewer losses (Wilcox signed rank test V = 48, p = 0.037). There was no significant difference in wins (Wilcox signed rank test V = 21, p = 0.54) or losses (Wilcoxon signed rank test V = 16, p = 0.832) in non-rising subdominants on the day of sham removal when compared to the day before alpha removal (**Figure 2C)**. In the alpha removed group, latency to first win occurred on average at 14.9 minutes, with some individuals winning their first fight within 15 seconds. This was significantly different from in the sham removal group, where latency to first win occurred on average at 34.9 minutes (Wilcoxon rank sum test: W = 36.5, p = 0.042; **Figure 2D)**.

**Figure 2.**
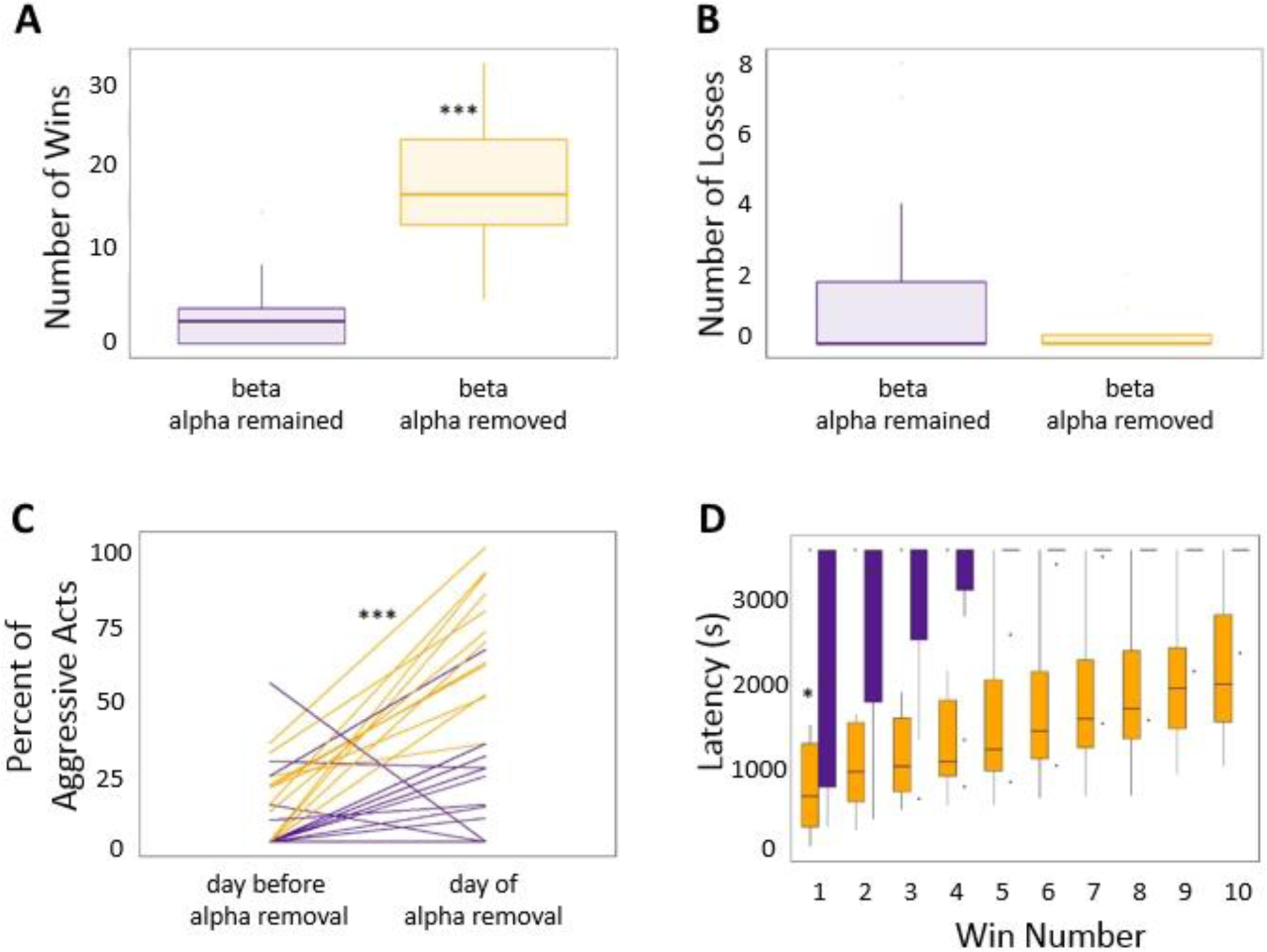
Behavioral changes in subdominant male following removal or non-removal of the alpha male. (A) Beta males in groups where the alpha male was removed win significantly more fights than beta males in groups where the alpha male remained in the group. (B) Beta males in groups where the alpha male was removed do not lose significantly fewer fights than beta males in groups where the alpha male was removed from the group. (C) The percentage of all contests won by the beta male on the day before removal or sham removal compared to the day after removal or sham removal. Lines represent different individuals. (D) Latency of subdominant males to win successive contests within one hour after the alpha male was removed (yellow) or shamremoved (purple)

In 11/12 of the removals, the rising subdominant was predicted based on data from the previous three days prior to the alpha removal. In these 11 cases, the rising subdominant was the male with the second highest Glicko ranking (i.e. the beta male) prior to removal. In the one instance where this was not the case, the individual that ascended had a slightly lower Glicko ranking than the previous beta male. However, it is worth noting that this was in the sixth removal after many manipulations of the social group, and the alpha male in this group was extremely despotic performing over 80% of all aggressive acts within the group. Consequently, fewer social contests occurred between lower-ranked individuals making it difficult to unequivocally identify the ranks of all other lower-ranked males at this time-point.

### Social ascent is associated with differential Fos immunoreactivity throughout the brain

**Table 1** describes Fos immunoreactivity pattern for beta males in the alpha removed and alpha remained conditions for each brain region studied. For 15/25 brain regions, there was a significant difference in Fos immunoreactivity, with individuals from the alpha removed group displaying significantly higher numbers of immunoreactive cells **(Table 1).** Consistent with our predictions, 6 of these regions (BNST, LS, AH, mPOA, dlPAG, vlPAG) are areas within the Social Behavior Network. The remaining 9 regions included prefrontal cortex (cingulate, infralimbic, prelimbic) as well as the retrosplenial cortex, hippocampal regions (CA1 and dentate gyrus), and a hypothalamic region (arcuate nucleus). Both the auditory cortex and visual cortex displayed increased immunoreactivity in the alpha removed condition. See **Figure 3** for sample images of Fos staining.

**Table 1.**
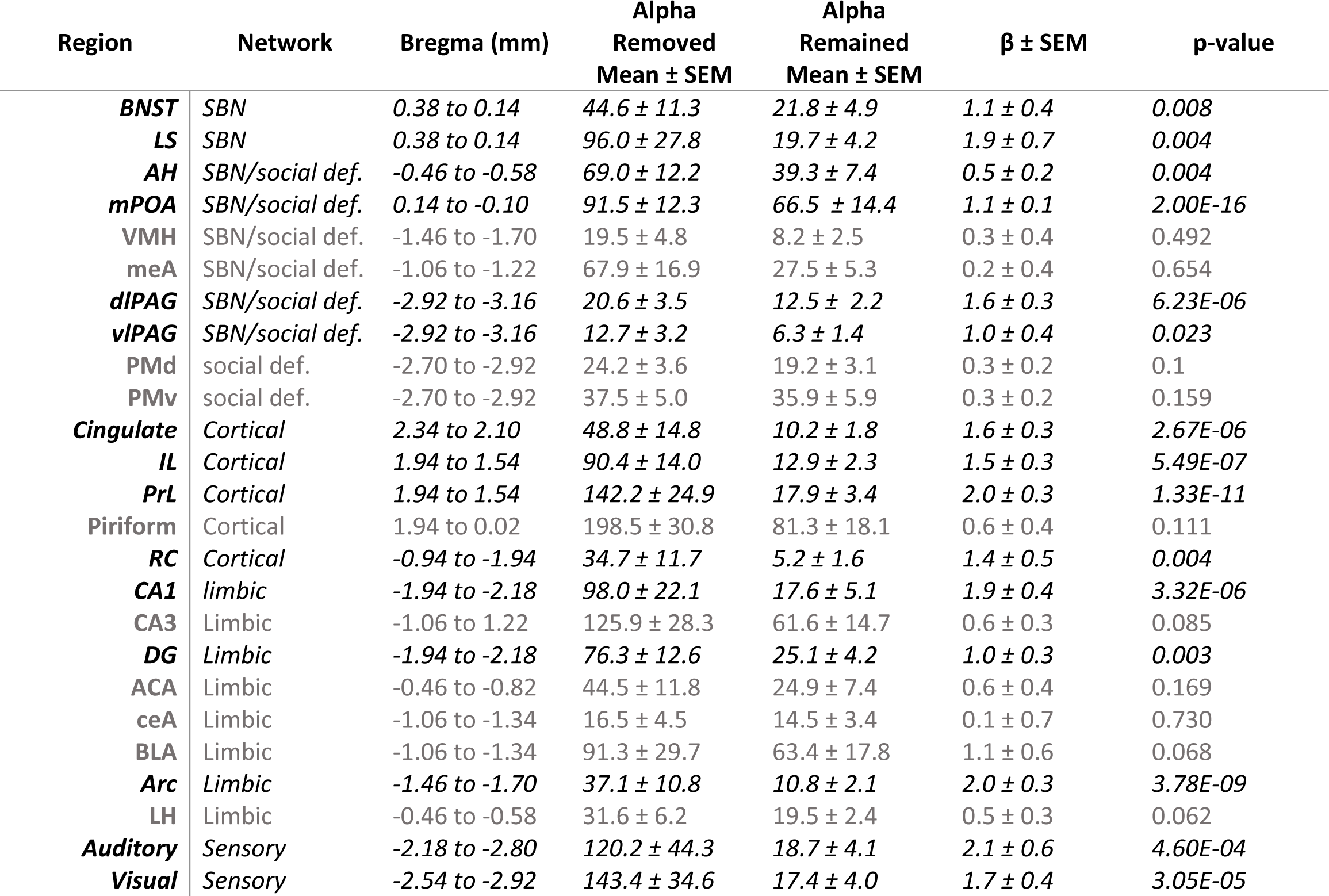
Table of Fos cell counts and results of mixed models for all brain regions examined.

| Region | Network | Bregma (mm) | Alpha<br>Removed<br>Mean ± SEM | Alpha<br>Remained<br>Mean ± SEM | $\beta \pm$ SEM | p-value |
| --- | --- | --- | --- | --- | --- | --- |
| **BNST** | *SBN* | *0.38 to 0.14* | *44.6 ± 11.3* | *21.8 ± 4.9* | *1.1 ± 0.4* | *0.008* |
| **LS** | *SBN* | *0.38 to 0.14* | *96.0 ± 27.8* | *19.7 ± 4.2* | *1.9 ± 0.7* | *0.004* |
| **AH** | *SBN/social def.* | *-0.46 to -0.58* | *69.0 ± 12.2* | *39.3 ± 7.4* | *0.5 ± 0.2* | *0.004* |
| **mPOA** | *SBN/social def.* | *0.14 to -0.10* | *91.5 ± 12.3* | *66.5 ± 14.4* | *1.1 ± 0.1* | *2.00E-16* |
| **VMH** | *SBN/social def.* | *-1.46 to -1.70* | *19.5 ± 4.8* | *8.2 ± 2.5* | *0.3 ± 0.4* | *0.492* |
| **meA** | *SBN/social def.* | *-1.06 to -1.22* | *67.9 ± 16.9* | *27.5 ± 5.3* | *0.2 ± 0.4* | *0.654* |
| **dIPAG** | *SBN/social def.* | *-2.92 to -3.16* | *20.6 ± 3.5* | *12.5 ± 2.2* | *1.6 ± 0.3* | *6.23E-06* |
| **vIPAG** | *SBN/social def.* | *-2.92 to -3.16* | *12.7 ± 3.2* | *6.3 ± 1.4* | *1.0 ± 0.4* | *0.023* |
| **PMd** | *social def.* | *-2.70 to -2.92* | *24.2 ± 3.6* | *19.2 ± 3.1* | *0.3 ± 0.2* | *0.1* |
| **PMv** | *social def.* | *-2.70 to -2.92* | *37.5 ± 5.0* | *35.9 ± 5.9* | *0.3 ± 0.2* | *0.159* |
| **Cingulate** | *Cortical* | *2.34 to 2.10* | *48.8 ± 14.8* | *10.2 ± 1.8* | *1.6 ± 0.3* | *2.67E-06* |
| **IL** | *Cortical* | *1.94 to 1.54* | *90.4 ± 14.0* | *12.9 ± 2.3* | *1.5 ± 0.3* | *5.49E-07* |
| **PrL** | *Cortical* | *1.94 to 1.54* | *142.2 ± 24.9* | *17.9 ± 3.4* | *2.0 ± 0.3* | *1.33E-11* |
| **Piriform** | *Cortical* | *1.94 to 0.02* | *198.5 ± 30.8* | *81.3 ± 18.1* | *0.6 ± 0.4* | *0.111* |
| **RC** | *Cortical* | *-0.94 to -1.94* | *34.7 ± 11.7* | *5.2 ± 1.6* | *1.4 ± 0.5* | *0.004* |
| **CA1** | *limbic* | *-1.94 to -2.18* | *98.0 ± 22.1* | *17.6 ± 5.1* | *1.9 ± 0.4* | *3.32E-06* |
| **CA3** | *Limbic* | *-1.06 to 1.22* | *125.9 ± 28.3* | *61.6 ± 14.7* | *0.6 ± 0.3* | *0.085* |
| **DG** | *Limbic* | *-1.94 to -2.18* | *76.3 ± 12.6* | *25.1 ± 4.2* | *1.0 ± 0.3* | *0.003* |
| **ACA** | *Limbic* | *-0.46 to -0.82* | *44.5 ± 11.8* | *24.9 ± 7.4* | *0.6 ± 0.4* | *0.169* |
| **ceA** | *Limbic* | *-1.06 to -1.34* | *16.5 ± 4.5* | *14.5 ± 3.4* | *0.1 ± 0.7* | *0.730* |
| **BLA** | *Limbic* | *-1.06 to -1.34* | *91.3 ± 29.7* | *63.4 ± 17.8* | *1.1 ± 0.6* | *0.068* |
| **Arc** | *Limbic* | *-1.46 to -1.70* | *37.1 ± 10.8* | *10.8 ± 2.1* | *2.0 ± 0.3* | *3.78E-09* |
| **LH** | *Limbic* | *-0.46 to -0.58* | *31.6 ± 6.2* | *19.5 ± 2.4* | *0.5 ± 0.3* | *0.062* |
| **Auditory** | *Sensory* | *-2.18 to -2.80* | *120.2 ± 44.3* | *18.7 ± 4.1* | *2.1 ± 0.6* | *4.60E-04* |
| **Visual** | *Sensory* | *-2.54 to -2.92* | *143.4 ± 34.6* | *17.4 ± 4.0* | *1.7 ± 0.4* | *3.05E-05* |

**Figure 3.**
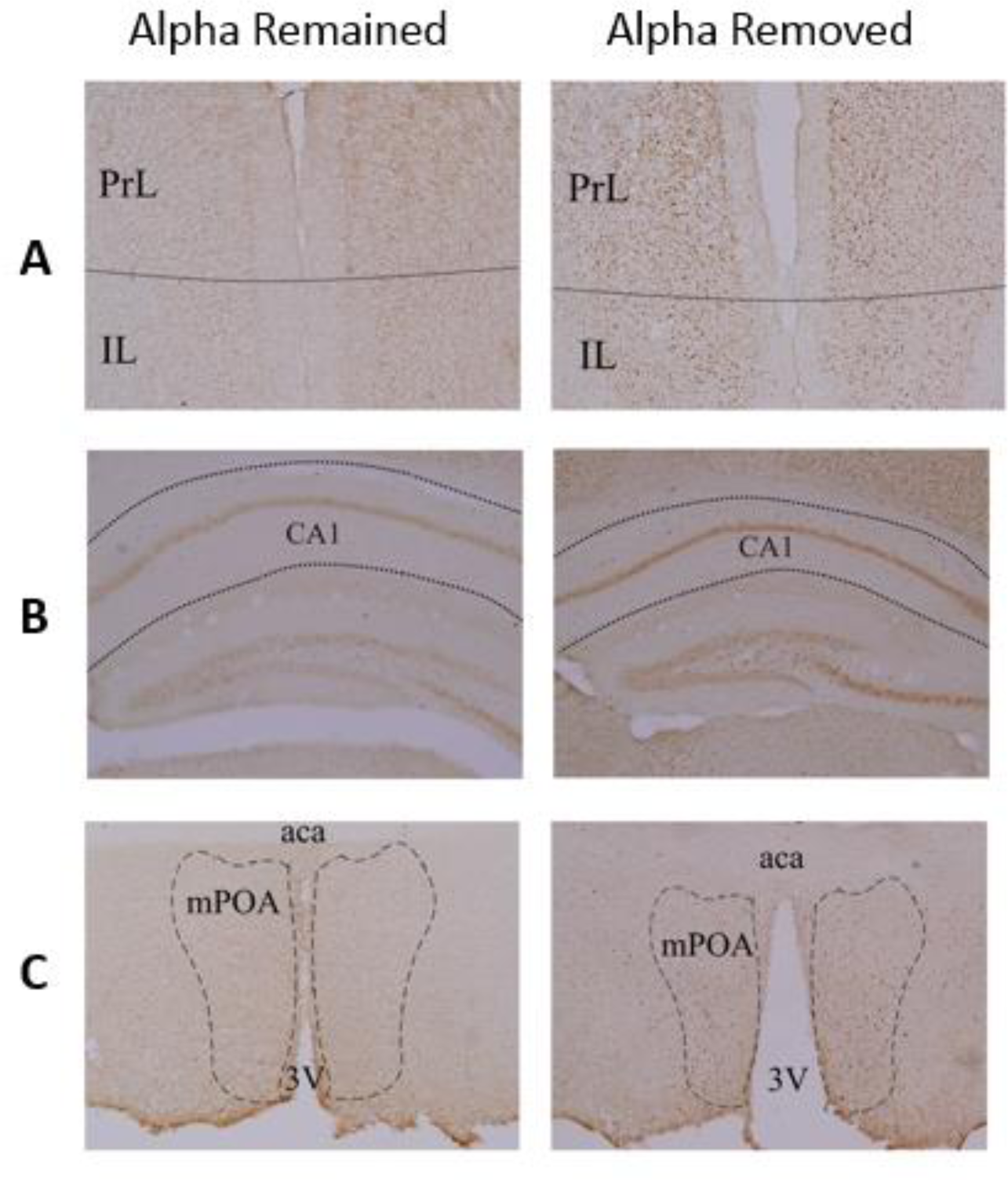
Sample images of Fos staining in individuals from the alpha remained (left) and alpha removed (right) groups. (A) Prelimbic and infralimbic regions of the medial prefrontal cortex (B) CA1 region of the hippocampus (C) Medial preoptic area of the hypothalamus

### Hierarchical clustering analysis suggests differential patterns of activation in individuals undergoing social ascent

To examine whether social ascent lead to differential co-activation patterns throughout the brain, we performed a hierarchical clustering analysis. Significantly different clusters between individuals undergoing social ascent and those in stable social groups were identified. In the alpha removed group, two distinct clusters formed, one including IL, PrL, DG, LS, CA3, vlPAG, LH, meA, Cing, RC, BLA, BNST, and CA1 and one including Aud, AH, PMv, CeA, Vis, CortA, dlPAG, mPOA, Pir, PMd, ARC, and VMH **(Figure 4A).** In the alpha remained group one cluster contained all regions but the BLA, IL, and PrL, with the IL and PrL splitting off into their own cluster **(Figure 4B)**. Notably, in the alpha removed group we saw greater positive correlation between brain regions **(Figure 4C)** and in the alpha remained group, we saw greater negative correlation between brain regions **(Figure 4D)**, suggesting that there was generally more coordinated activation of pathways in the individuals undergoing social ascent.

**Figure 4.**
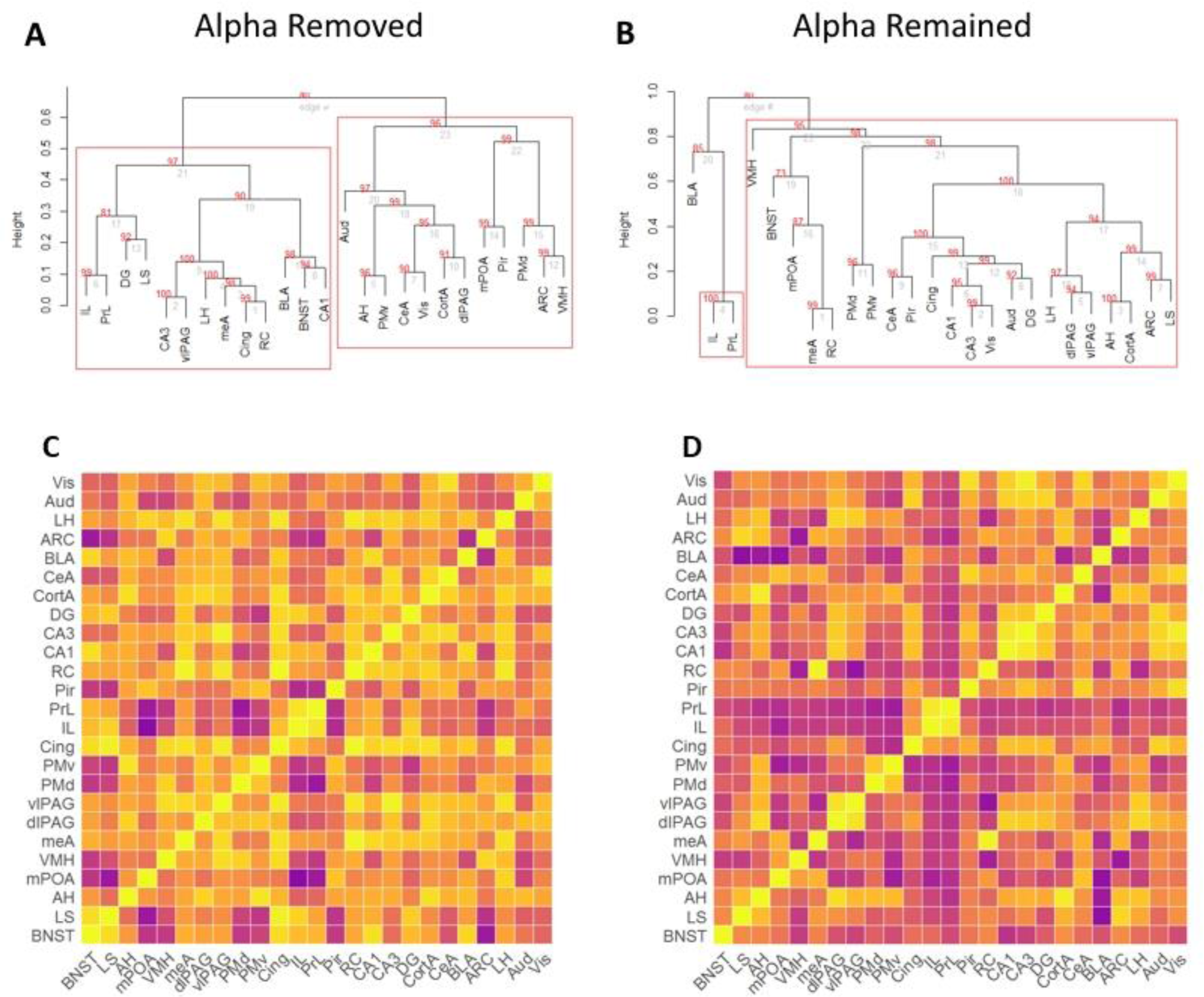
Beta individuals undergoing social ascent have distinct neural activation patterns as compared to beta individuals from stable groups. (A and B) Cluster dendrogram displaying results of hierarchical clustering analysis for beta males where the alpha was removed (A) and for beta males where the alpha remained (B). Red numbers indicate the approximately unbiased p-value generated through multiscale bootstrap resampling. Values higher than 95 indicate statistical significance. Red boxes denote significant clusters that are strongly supported by data. (C and D) Pearson correlation coefficients were used to create a heatmap of neural co-activation across examined brain regions. (C) Heatmap for beta males where the alpha was removed (D) Heatmap for beta males where the alpha remained

## DISCUSSION

In the present study, we successfully replicated the behavioral findings from our previous work (Williamson, Romeo, et al., 2017), illustrating that following removal of the alpha male, beta males recognized the emergence of a power vacuum and use this opportunity to ascend to alpha status. Ascending males won significantly more and lost significantly less in comparison to their own behavior the day before, as well as compared to non-ascending beta males in hierarchies whose alpha had not been removed. Moreover, this change occurs rapidly with beta males beginning their ascent on average within 15 minutes. These findings provide further evidence that individuals in social groups recognize and behaviorally respond to dynamic changes in social context. This ability appears to be a fundamental feature of living within a social group, and has been seen to occur in a similarly controlled manner in African cichlid fish (Maruska, Zhang, et. al., 2013; Maruska & Fernald, 2010) where beta males begin to change color and increase aggression in response to alpha removal within minutes, and in primates, where beta males quickly and forcefully ascended to alpha status following alpha males receiving amygdaloid lesions (Rosvold, Mirsky, & Pribram, 1954). This ability for individuals to recognize and rapidly respond to changes in social status is an essential feature of social competence and is associated with greater social, reproductive, and health outcomes (Hofmann et al., 2014; Taborsky & Oliveira, 2012).

In the current study, we also demonstrate that response to a change in social context is associated increases in neural activity. Notable increases in immediate early immunoreactivity were observed throughout the SBN as well as in the prefrontal cortex and hippocampus. It appears that coordinated activation of these regions is required to facilitate the assessment of a change in the social context combined with the increase in aggressive behavior to facilitate social ascent. In the SBN, we saw significant differences in cell counts between ascending beta males and stable beta males in the BNST, lateral septum, mPOA, anterior hypothalamus, and both the dlPAG and vlPAG. Each of these regions have well-established roles in the modulation of social behavior (Goodson, 2005). Significantly, we did not see any difference in neural activity in the VMH or medial amygdala. The VMH has been demonstrated to be of particular interest in female social behavior (Goodson, 2005), for example when females are assessing the social dominance of potential mates (Desjardins, Klausner, & Fernald, 2010) or in female aggression (Hashikawa et al., 2017). Further, the VMH has been shown to be involved in response to territorial challenge and stress (Goodson & Evans, 2004) but does not appear to be involved in the processing of social context information. The lack of difference in the medial amygdala is more remarkable, as it has been heavily implicated in social dominance (Bolhuis, Fitzgerald, et. al., 1984; Rosvold, Enger et al., 1954; So et al., 2015; Timmer, Cordero, et. al., 2011). However, these previous findings appear to be specific to stable social groups and to understanding the physiology of individuals of dominant vs. subordinate status and cannot be assumed to extend to individuals responding to a changing social context and subsequently undergoing a change in social status.

The largest brain differences observed between ascending beta males and non-ascending beta males were in the prelimbic and infralimbic regions of the prefronal cortex. In non-ascending beta males we find very little activation in these regions, whereas in ascending males we find very large levels of activation. These regions of the medial prefrontal cortex have been established as essential to the processing of rodent social behavior, including aggressive behavior, affiliative behaviors, and dominance behavior (Ko, 2017; Wang, Kessels, & Hu, 2014). In mice, prelimbic neurons have also been implicated in processing social preference as well as social-spatial information (Murugan et al., 2017). Further, the mPFC has been implicated in humans in the processing of social status information (Silk et al., 2017; Wang et al., 2014; Zerubavel et al., 2015) and the social network position of others (Parkinson, Kleinbaum, & Wheatley, 2017) as well as processing information in relation to self – i.e. how one fits into the broader social context (Pfeifer et al., 2009). Other studies have shown activation of the mPFC when processing unstable social hierarchies (Zink et al., 2008). In pairs of rhesus monkeys, the dominant individual’s PFC becomes locked in an “up-state” while the subordinate individual’s becomes locked in a “down-state”. This state rapidly switches when relative hierarchical status is switched (Fujii, Hihara, Nagasaka, & Iriki, 2009). Our findings provide further evidence for the crucial role of the medial prefrontal cortex in processing information about social context. Moreover, we purposefully chose to examine brains 90 minutes after each male had behaviorally demonstrated that they had begun to socially ascend. The aim of this methodological approach was to ensure we could identify regions involved in processing the change in social context and that might promote further downstream brain activation. Taking into account the previous literature and our current findings, we suggest that the IL and PL are key regions in the tracking of changes in the social environment and in facilitating further social-decision making, i.e. social ascent.

Our analyses also identified neural activity within the hippocampus, specifically the CA1 and dentate gyrus, as being associated with dynamic change in social status. Neurons in the ventral hippocampus are necessary for social memory storage and are specifically activated in response to familiar mice (Okuyama, Kitamura, Roy, Itohara, & Tonegawa, 2016). Before a beta individual begins his ascent, we observe patrolling and olfactory exploration of the vivarium by these males. During this exploration, the beta male is coming into contact with the urine of the familiar alpha male, a highly salient signal of social status (Lee, Khan, & Curley, 2017). This exploration could potentially lead to activation of the CA1 in response to social memory of interaction with the alpha. This social information could be integrated with the corresponding excitation in the prefrontal cortices. Notably, there is a clear, excitatory pathway from CA1 to the prelimbic region of the prefrontal cortex (Thierry, Gioanni, Dégénétais, & Glowinski, 2000), though further studies are required to elucidate the exact activation patterns connecting these brain regions. Further, both the dentate gyrus and the retropslenial cortex have been implicated in regulating spatial memory (Ibi et al., 2008; Jessberger et al., 2009; Nilsson, Perfilieva, Johansson, Orwar, & S. Eriksson, 1999; Ophir, Wolff, & Phelps, 2008; Czajkowski et al., 2014). In a changing complex social environment, determining the physical presence or absence of more dominant individuals is critical and the observed activation of these brain regions in ascending males may be related to utilization of spatial memory. Moreover, oxytocin receptor signaling in the dentate gyrus has also been shown to be necessary for discrimination of social stimuli (Raam, McAvoy, Besnard, Veenema, & Sahay, 2017), providing further evidence for the importance of this brain region in processing social contextual cues.

We observed significantly elevated activation in the primary visual cortex and primary auditory cortex of ascending males. While studies of mouse social behavior often do not focus on the primary visual cortex, it is essential to social processing in humans – social visual signals provide information about emotional expression, direction of gaze, body posture, and movement, all important social cues (Adolphs, 2003). Studies in non-human primates have demonstrated that neuronal responses in the visual cortex appear to encode highly specific social stimuli such as those described above (i.e. faces, gaze, etc.) (Perrett, Rolls, & Caan, 1982). While there is limited work to suggest that processing of visual stimuli is essential to mouse social behavior, studies of mouse models of autism suggest that excitatory/inhibitory balance and plasticity in the visual cortex during critical periods in development is important for the development of social behavior (Gogolla et al., 2009). Further, lack of proper gamma oscillations generated in the primary visual cortex are similarly implicated in autism, and have been shown to be important for information processing and learning, suggesting that these oscillations are important for appropriate social behavior (Gogolla et al., 2009; Singer, 1993). It is likely that the activation of the visual cortex during social opportunity is related to visual monitoring of the social environment. The impact of changing social context on the primary auditory cortex is consistent with the established role of auditory cues in communication in mice. Ultrasonic vocalizations (USVs) have been demonstrated to facilitate social interactions in mice (Liu, Miller, Merzenich, & Schreiner, 2003). Dominant males have been shown to elicit significantly more of these vocalizations in mating situations (Lumley, Sipos, Charles, Charles, & Meyerhoff, 1999; Nyby, Dizinno, & Whitney, 1976), though not necessarily during aggressive encounters (Nyby & Whitney, 1978; Portfors, 2007). These USVs may function as territorial signals between males mice (Gourbal, Barthelemy, Petit, & Gabrion, 2004; Hammerschmidt, Radyushkin, Ehrenreich, & Fischer, 2012). These findings lead us to hypothesize that alpha males most likely emit USVs on a regular basis in the vivarium and that subdominants process the auditory inputs from the environment to determine if the alpha male is present or absent.

Our analyses of immediate early gene activation indicated different co-activation patterns in the beta males undergoing social ascent from those in a stable social group. Most notably, those in the alpha removed group had overall increased, positively correlated activation throughout the regions studied, where those in the alpha remained group showed a more negative correlation throughout these regions. This is consistent with the idea that individuals whose social context is rapidly changing are responding with a cascade of neural activation to coordinate complex social behavior. Further, we show through a hierarchical clustering analysis that individuals in the alpha removed group have more specific clusters of activation, with prefrontal cortical (IL, PL, cingulate and retrosplenial cortices), hippocampal (CA1, CA3 and dentate gyrus), and some SBN regions clustering separately from sensory regions, premammillary nuclei, periaqueductal grey, and other hypothalamic regions including the mPOA. In non-ascending males, the IL and PL cortices are in a separate cluster altogether from the SBN. Guided by these data, we propose that when a social opportunity arises, the IL and PL cortices are activated and evaluate changes in the social context. This information is integrated along with other information related to social and spatial memory processed by the hippocampus and retrosplenial cortex in a coordinated fashion. Information from sensory regions is synchronously activated and integrated with this social information before activating nodes of the SBN to facilitate the appropriate behavioral response.

## CONCLUSION

In the present study, we established that following removal of the alpha male from a stable dominance hierarchy, the beta male recognizes the absence of the alpha and responds by rapidly increasing aggressive behaviors. We have demonstrated that this salient social stimulus of removing the alpha male and subsequent change in behavior by the beta male leads to increased Fos immunoreactivity throughout the brain, specifically in regions of the SBN, as well as the medial prefrontal cortex, retrosplenial cortex and area CA1 of the hippocampus. These findings suggest that the complex social competence required to assess one’s social context and respond appropriately is modulated by a synchronous and integrated increase in activity throughout the brain.

## ACKNOWLEDGMENTS

We would like to thank all members of the Curley Lab for their help in collecting behavioral data as well as Alesi Lanham for her assistance with the immunohistochemistry.

## REFERENCES

Adolphs, R. (2003). Cognitive neuroscience: Cognitive neuroscience of human social behaviour. Nature Reviews Neuroscience, 4(3), 165. https://doi.org/10.1038/nrn1056

Bates, D., Maechler, M., Bolker, B., Walker, S., Christensen, R. H. B., Singmann, H., … Grothendieck, G. (2015). lme4: Linear Mixed-Effects Models using “Eigen” and S4 (Version 1.1-10). Retrieved from https://cran.r-project.org/web/packages/lme4/index.html

Bolhuis, J. J., Fitzgerald, R. E., Dijk, D. J., & Koolhaas, J. M. (1984). The corticomedial amygdala and learning in an agonistic situation in the rat. Physiology & Behavior, 32(4), 575–579. https://doi.org/10.1016/0031-9384(84)90311-1

Burmeister, S. S., Jarvis, E. D., & Fernald, R. D. (2005). Rapid Behavioral and Genomic Responses to Social Opportunity. PLOS Biol, 3(11), e363. https://doi.org/10.1371/journal.pbio.0030363

Chase, I. D., & Seitz, K. (2011). Self-structuring properties of dominance hierarchies: A new perspective. Advances in Genetics, 75, 51.

Chase, I. D., Tovey, C., Spangler-Martin, D., & Manfredonia, M. (2002). Individual differences versus social dynamics in the formation of animal dominance hierarchies. Proceedings of the National Academy of Sciences, 99(8), 5744–5749.

Curley, J. P. (2016a). compete: Organizing and Analyzing Social Dominance Hierarchy Data. (Version 0.1). Retrieved from https://cran.r-project.org/web/packages/compete/index.html

Curley, J. P. (2016b). Temporal pairwise-correlation analysis provides empirical support for attention hierarchies in mice. Biology Letters, 12(5), 20160192. https://doi.org/10.1098/rsbl.2016.0192

Czajkowski, R., Jayaprakash, B., Wiltgen, B., Rogerson, T., Guzman-Karlsson, M. C., Barth, A. L., … Silva, A. J. (2014). Encoding and storage of spatial information in the retrosplenial cortex. Proceedings of the National Academy of Sciences, 111(23), 8661–8666. https://doi.org/10.1073/pnas.1313222111

De Vries, H. (1995). An improved test of linearity in dominance hierarchies containing unknown or tied relationships. Animal Behaviour, 50(5), 1375–1389.

Desjardins, J. K., Klausner, J. Q., & Fernald, R. D. (2010). Female genomic response to mate information. Proceedings of the National Academy of Sciences, 107(49), 21176–21180. https://doi.org/10.1073/pnas.1010442107

Fernald, R. D. (2014). Cognitive Skills Needed for Social Hierarchies. Cold Spring Harbor Symposia on Quantitative Biology, 79, 229–236. https://doi.org/10.1101/sqb.2014.79.024752

Fujii, N., Hihara, S., Nagasaka, Y., & Iriki, A. (2009). Social state representation in prefrontal cortex. Social Neuroscience, 4(1), 73–84. https://doi.org/10.1080/17470910802046230

Glickman, M. E. (1999). Parameter estimation in large dynamic paired comparison experiments. Journal of the Royal Statistical Society: Series C (Applied Statistics), 48(3), 377–394.

Gogolla, N., LeBlanc, J. J., Quast, K. B., Südhof, T. C., Fagiolini, M., & Hensch, T. K. (2009). Common circuit defect of excitatory-inhibitory balance in mouse models of autism. Journal of Neurodevelopmental Disorders, 1(2), 172. https://doi.org/10.1007/s11689-009-9023-x

Goodson, J. L. (2005). The Vertebrate Social Behavior Network: Evolutionary Themes and Variations. Hormones and Behavior, 48(1), 11–22. https://doi.org/10.1016/j.yhbeh.2005.02.003

Goodson, J. L., & Evans, A. K. (2004). Neural responses to territorial challenge and nonsocial stress in male song sparrows: segregation, integration, and modulation by a vasopressin V1 antagonist. Hormones and Behavior, 46(4), 371–381. https://doi.org/10.1016/j.yhbeh.2004.02.008

Gourbal, B. E. F., Barthelemy, M., Petit, G., & Gabrion, C. (2004). Spectrographic analysis of the ultrasonic vocalisations of adult male and female BALB/c mice. Naturwissenschaften, 91(8), 381–385. https://doi.org/10.1007/s00114-004-0543-7

Grosenick, L., Clement, T. S., & Fernald, R. D. (2007). Fish can infer social rank by observation alone. Nature, 445(7126), 429–432.

Hammerschmidt, K., Radyushkin, K., Ehrenreich, H., & Fischer, J. (2012). The Structure and Usage of Female and Male Mouse Ultrasonic Vocalizations Reveal only Minor Differences. PLOS ONE, 7(7), e41133. https://doi.org/10.1371/journal.pone.0041133

Hashikawa, K., Hashikawa, Y., Tremblay, R., Zhang, J., Feng, J. E., Sabol, A., … Lin, D. (2017). Esr1+ cells in the ventromedial hypothalamus control female aggression. Nature Neuroscience, advance online publication. https://doi.org/10.1038/nn.4644

Hofmann, H. A., Beery, A. K., Blumstein, D. T., Couzin, I. D., Earley, R. L., Hayes, L. D., … Rubenstein, D. R. (2014). An evolutionary framework for studying mechanisms of social behavior. Trends in Ecology & Evolution, 29(10), 581–589. https://doi.org/10.1016/j.tree.2014.07.008

Holmes, M. M., Goldman, B. D., & Forger, N. G. (2008). Social status and sex independently influence androgen receptor expression in the eusocial naked mole-rat brain. Hormones and Behavior, 54(2), 278–285. https://doi.org/10.1016/j.yhbeh.2008.03.010

Huffman, L. S., Hinz, F. I., Wojcik, S., Aubin-Horth, N., & Hofmann, H. A. (2015). Arginine vasotocin regulates social ascent in the African cichlid fish Astatotilapia burtoni. General and Comparative Endocrinology, 212, 106–113. https://doi.org/10.1016/j.ygcen.2014.03.004

Ibi, D., Takuma, K., Koike, H., Mizoguchi, H., Tsuritani, K., Kuwahara, Y., … Yamada, K. (2008). Social isolation rearing-induced impairment of the hippocampal neurogenesis is associated with deficits in spatial memory and emotion-related behaviors in juvenile mice. Journal of Neurochemistry, 105(3), 921–932. https://doi.org/10.1111/j.1471-4159.2007.05207.x

Jessberger, S., Clark, R. E., Broadbent, N. J., Clemenson, G. D., Consiglio, A., Lie, D. C., … Gage, F. H. (2009). Dentate gyrus-specific knockdown of adult neurogenesis impairs spatial and object recognition memory in adult rats. Learning & Memory, 16(2), 147–154. https://doi.org/10.1101/lm.1172609

Ko, J. (2017). Neuroanatomical Substrates of Rodent Social Behavior: The Medial Prefrontal Cortex and Its Projection Patterns. Frontiers in Neural Circuits, 11. https://doi.org/10.3389/fncir.2017.00041

Kucharski, R., Maleszka, J., Foret, S., & Maleszka, R. (2008). Nutritional Control of Reproductive Status in Honeybees via DNA Methylation. Science, 319(5871), 1827–1830. https://doi.org/10.1126/science.1153069

Lee, W., Khan, A., & Curley, J. P. (2017). Major urinary protein levels are associated with social status and context in mouse social hierarchies. Proc. R. Soc. B, 284(1863), 20171570. https://doi.org/10.1098/rspb.2017.1570

Lein, E. S., Hawrylycz, M. J., Ao, N., Ayres, M., Bensinger, A., Bernard, A., … Jones, A. R. (2007). Genome-wide atlas of gene expression in the adult mouse brain. Nature, 445(7124), 168–176. https://doi.org/10.1038/nature05453

Liu, R. C., Miller, K. D., Merzenich, M. M., & Schreiner, C. E. (2003). Acoustic variability and distinguishability among mouse ultrasound vocalizations. The Journal of the Acoustical Society of America, 114(6), 3412–3422. https://doi.org/10.1121/1.1623787

Maruska, K. P., Zhang, A., Neboori, A., & Fernald, R. D. (2013). Social Opportunity Causes Rapid Transcriptional Changes in the Social Behaviour Network of the Brain in an African Cichlid Fish. Journal of Neuroendocrinology, 25(2), 145–157. https://doi.org/10.1111/j.1365-2826.2012.02382.x

Lumley, L. A., Sipos, M. L., Charles, R. C., Charles, R. F., & Meyerhoff, J. L. (1999). Social Stress Effects on Territorial Marking and Ultrasonic Vocalizations in Mice. Physiology & Behavior, 67(5), 769–775. https://doi.org/10.1016/S0031-9384(99)00131-6

Maruska, Karen P., & Fernald, R. D. (2010). Behavioral and physiological plasticity: Rapid changes during social ascent in an African cichlid fish. Hormones and Behavior, 58(2), 230–240. https://doi.org/10.1016/j.yhbeh.2010.03.011

Muller, M. N., & Wrangham, R. W. (2004). Dominance, aggression and testosterone in wild chimpanzees: a test of the “challenge hypothesis.” *Animal Behaviour*, 67(1), 113–123. https://doi.org/10.1016/j.anbehav.2003.03.013

Murugan, M., Jang, H. J., Park, M., Miller, E. M., Cox, J., Taliaferro, J. P., … Witten, I. B. (2017). Combined Social and Spatial Coding in a Descending Projection from the Prefrontal Cortex. Cell, 0(0). https://doi.org/10.1016/j.cell.2017.11.002

Newman, S. W. (1999). The Medial Extended Amygdala in Male Reproductive Behavior A Node in the Mammalian Social Behavior Network. Annals of the New York Academy of Sciences, 877(1), 242–257. https://doi.org/10.1111/j.1749-6632.1999.tb09271.x

Nilsson, M., Perfilieva, E., Johansson, U., Orwar, O., & S. Eriksson, P. (1999). Enriched environment increases neurogenesis in the adult rat dentate gyrus and improves. Journal of Neurobiology, 39, 569–578. https://doi.org/10.1002/(SICI)1097-4695(19990615)39:4<569::AID-NEU10>3.0.CO;2- F

Noonan, M. P., Sallet, J., Mars, R. B., Neubert, F. X., O’Reilly, J. X., Andersson, J. L., … Rushworth, M. F. S. (2014). A Neural Circuit Covarying with Social Hierarchy in Macaques. PLOS Biology, 12(9), e1001940. https://doi.org/10.1371/journal.pbio.1001940

Nyby, J., Dizinno, G. A., & Whitney, G. (1976). Social status and ultrasonic vocalizations of male mice. Behavioral Biology, 18(2), 285–289. https://doi.org/10.1016/S0091-6773(76)92198-2

Nyby, J., & Whitney, G. (1978). Ultrasonic communication of adult myomorph rodents. Neuroscience & Biobehavioral Reviews, 2(1), 1–14. https://doi.org/10.1016/0149--7634(78)90003-9

O’Connell, L. A., & Hofmann, H. A. (2011). The Vertebrate mesolimbic reward system and social behavior network: A comparative synthesis. The Journal of Comparative Neurology, 519(18), 3599–3639. https://doi.org/10.1002/cne.22735

Okuyama, T., Kitamura, T., Roy, D. S., Itohara, S., & Tonegawa, S. (2016). Ventral CA1 neurons store social memory. Science, 353(6307), 1536–1541. https://doi.org/10.1126/science.aaf7003

Oliveira, R. F. (2009). Social behavior in context: Hormonal modulation of behavioral plasticity and social competence. Integrative and Comparative Biology, 49(4), 423–440. https://doi.org/10.1093/icb/icp055

Oliveira, R. F., & Almada, V. C. (1996). On the (in)stability of dominance hierarchies in the cichlid fish Oreochromis mossambicus. Aggressive Behavior, 22(1), 37–45. https://doi.org/10.1002/(SICI)1098-2337(1996)22:1<37::AID-AB4>3.0.CO;2-R

Ophir, A. G., Wolff, J. O., & Phelps, S. M. (2008). Variation in neural V1aR predicts sexual fidelity and space use among male prairie voles in semi-natural settings. Proceedings of the National Academy of Sciences, 105(4), 1249–1254. https://doi.org/10.1073/pnas.0709116105

Parkinson, C., Kleinbaum, A. M., & Wheatley, T. (2017). Spontaneous neural encoding of social network position. Nature Human Behaviour, 1(5), 0072. https://doi.org/10.1038/s41562-017-0072

Paxinos, G., & Franklin, K. B. J. (2004). The Mouse Brain in Stereotaxic Coordinates. Gulf Professional Publishing.

Perrett, D. I., Rolls, E. T., & Caan, W. (1982). Visual neurones responsive to faces in the monkey temporal cortex. Experimental Brain Research, 47(3), 329–342. https://doi.org/10.1007/BF00239352

Pfeifer, J. H., Masten, C. L., Borofsky, L. A., Dapretto, M., Fuligni, A. J., & Lieberman, M. D. (2009). Neural Correlates of Direct and Reflected Self-Appraisals in Adolescents and Adults: When Social Perspective-Taking Informs Self-Perception. Child Development, 80(4), 1016–1038. https://doi.org/10.1111/j.1467-8624.2009.01314.x

Portfors, C. V. (2007). Types and Functions of Ultrasonic Vocalizations in Laboratory Rats and Mice. Journal of the American Association for Laboratory Animal Science, 46(1), 28–34.

R Core Team. (2016). R: A language and environment for statistical computing (Version 3.2.3). Vienna, Austria: R Foundation for Statistical Computing. Retrieved from https://www.R-project.org/

Raam, T., McAvoy, K. M., Besnard, A., Veenema, A., & Sahay, A. (2017). Hippocampal oxytocin receptors are necessary for discrimination of social stimuli. Nature Communications, 8(1), 2001. https://doi.org/10.1038/s41467-017-02173-0

Rosvold, H. Enger, H., Mirsky, A. F., & Pribram, K. H. (1954). Influence of amygdalectomy on social behavior in monkeys. Journal of Comparative and Physiological Psychology, 47(3), 173–178. https://doi.org/10.1037/h0058870

RStudio Team. (2015). RStudio: Integrated Development Environment for R. Boston, MA: RStudio, Inc. Retrieved from http://www.rstudio.com/

Sapolsky, R. M. (1993). The physiology of dominance in stable versus unstable social hierarchies. In W. A. Mason & S. P. Mendoza (Eds.), Primate social conflict (pp. 171–204). Albany, NY, US: State University of New York Press.

Sarkar, D. (2017). Lattice: Multivariate Data Visualization with R (Version 0.20-35). New York.

Schneider, C. A., Rasband, W. S., & Eliceiri, K. W. (2012). NIH Image to ImageJ: 25 years of image analysis. Nature Methods, 9(7), 671–675. https://doi.org/10.1038/nmeth.2089

Silk, J. S., Lee, K. H., Kerestes, R., Griffith, J. M., Dahl, R. E., & Ladouceur, C. D. (2017). “Loser” or “Popular”?: Neural response to social status words in adolescents with major depressive disorder. Developmental Cognitive Neuroscience, 28(Supplement C), 1–11. https://doi.org/10.1016/j.dcn.2017.09.005

Singer, W. (1993). Synchronization of Cortical Activity and its Putative Role in Information Processing and Learning. Annual Review of Physiology, 55(1), 349–374. https://doi.org/10.1146/annurev.ph.55.030193.002025

So, N., Franks, B., Lim, S., & Curley, J. P. (2015). A social network approach reveals associations between mouse social dominance and brain gene expression. PloS One, 10(7), e0134509. https://doi.org/10.1371/journal.pone.0134509

Stephenson, A., & Sonas, J. (2012). PlayerRatings: Dynamic updating methods for player ratings estimation (Version 1.0-0). Retrieved from {http://CRAN.R-project.org/package=PlayerRatings

Suzuki, R., & Simodaira, H. (2015). pvclust: Hierarchical Clustering with P-Values via Multiscale Bootstrap Resampling (Version 2.0).

Taborsky, B., & Oliveira, R. F. (2012). Social competence: an evolutionary approach. Trends in Ecology & Evolution, 27(12), 679–688. https://doi.org/10.1016/j.tree.2012.09.003

Thierry, A.-M., Gioanni, Y., Dégénétais, E., & Glowinski, J. (2000). Hippocampo-prefrontal cortex pathway: Anatomical and electrophysiological characteristics. Hippocampus, 10(4), 411–419. https://doi.org/10.1002/1098-1063(2000)10:4<411::AID-HIPO7>3.0.CO;2-A

Timmer, M., Cordero, M. I., Sevelinges, Y., & Sandi, C. (2011). Evidence for a Role of Oxytocin Receptors in the Long-Term Establishment of Dominance Hierarchies. Neuropsychopharmacology, 36(11), 2349–2356. https://doi.org/10.1038/npp.2011.125

Wang, F., Kessels, H. W., & Hu, H. (2014). The mouse that roared: neural mechanisms of social hierarchy. Trends in Neurosciences, 37(11), 674–682. https://doi.org/10.1016/j.tins.2014.07.005

Wang, F., Zhu, J., Zhu, H., Zhang, Q., Lin, Z., & Hu, H. (2011). Bidirectional Control of Social Hierarchy by Synaptic Efficacy in Medial Prefrontal Cortex. Science, 334(6056), 693–697. https://doi.org/10.1126/science.1209951

Williamson, C. M., Franks, B., & Curley, J. P. (2016). Mouse Social Network Dynamics and Community Structure are Associated with Plasticity-Related Brain Gene Expression. Frontiers in Behavioral Neuroscience, 10. https://doi.org/10.3389/fnbeh.2016.00152

Williamson, C. M., Lee, W., & Curley, J. P. (2016). Temporal dynamics of social hierarchy formation and maintenance in male mice. Animal Behaviour, 115, 259–272. https://doi.org/10.1016/j.anbehav.2016.03.004

Williamson, C. M., Lee, W., Romeo, R. D., & Curley, J. P. (2017). Social context-dependent relationships between mouse dominance rank and plasma hormone levels. Physiology & Behavior, 171, 110–119. https://doi.org/10.1016/j.physbeh.2016.12.038

Williamson, C. M., Romeo, R. D., & Curley, J. P. (2017). Dynamic changes in social dominance and mPOA GnRH expression in male mice following social opportunity. Hormones and Behavior, 87, 80–88. https://doi.org/10.1016/j.yhbeh.2016.11.001

Zerubavel, N., Bearman, P. S., Weber, J., & Ochsner, K. N. (2015). Neural mechanisms tracking popularity in real-world social networks. Proceedings of the National Academy of Sciences, 112(49), 15072–15077. https://doi.org/10.1073/pnas.1511477112

Zink, C. F., Tong, Y., Chen, Q., Bassett, D. S., Stein, J. L., & Meyer-Lindenberg, A. (2008). Know Your Place: Neural Processing of Social Hierarchy in Humans. Neuron, 58(2), 273–283. https://doi.org/10.1016/j.neuron.2008.01.025

